# Ingestion of microplastic fibres, but not microplastic beads, impacts growth rates in the tropical house cricket *Gryllodes sigillatus*

**DOI:** 10.1101/2022.02.07.479438

**Authors:** Serita Fudlosid, Marshall W. Ritchie, Matthew J. Muzzatti, Jane E. Allison, Jennifer Provencher, Heath A. MacMillan

**Author notes:** **Correspondence:** Serita Fudlosid.

## Abstract

Microplastic is a growing concern as an environmental contaminant as it is ubiquitous in our ecosystems. Microplastics are present in terrestrial environments, yet the majority of studies have focused on the adverse effects of microplastics on aquatic biota. We hypothesized that microplastic ingestion by a terrestrial insect would have localized effects on gut health and nutrient absorption, such that prolonged dietary microplastic exposure would impact growth rate and adult body size. We further hypothesized that plastic form (fibres vs. beads) would influence these effects because of the nature of gut-plastic interactions. Freshly hatched tropical house crickets (*Gryllodes sigillatus*) were fed a standard diet containing different concentrations of either fluorescent polyethylene microplastic beads (75-105 μm), or untreated polyethylene terephthalate microfibers (<5 mm) until they died or reached adulthood (approximately 8 weeks). Weight and body length were measured weekly and microplastic ingestion was confirmed through fluorescence microscopy and visual inspection of the frass. While, to our surprise, we found no effect of polyethylene bead ingestion on growth rate or final body size of *G. sigillatus*, females experienced a reduction in size and weight when fed high concentrations of polyethylene terephthalate microfibers. These results suggest that high concentrations of polyethylene beads of the 100 μm size range can pass through the cricket gut without a substantial negative effect on their growth and development time, but high concentrations of polyethylene terephthalate microfibers cannot. Although we report the negative effects of microplastic ingestion on the growth of *G. sigillatus*, it remains uncertain what threats microplastics pose to terrestrial insects.

## Introduction

Plastic pollution is a major global environmental issue (Kumar et al., 2021). As of 2015, it was estimated that 6300 Mt of plastic waste had been produced since 1950 with only 9% recycled and 12% incinerated (Geyer et al., 2017). A combination of plastic’s persistence in the environment and limited waste recovery has attracted public attention to the risk posed to the environment and human health (Campanale et al., 2020; Chae & An, 2018; Ivar Do Sul & Costa, 2014). A major component of environmental plastic pollution is microplastic (MP); plastic particles <5 mm (Thompson et al., 2009). MPs are produced as either primary MPs (manufactured micro-sized plastic for commercial or domestic use), or secondary MPs (resulting from breakdown of larger plastics from mechanical fragmentation or photolytic/biological degradation) (Helmberger et al., 2020). Primary MPs in the environment can originate from sources such as pellet spillage from air blasting machines or microbeads in facial cleansers (Auta et al., 2017), whereas secondary MPs can have numerous sources, such as agricultural plastic films, synthetic rubber debris from tires, or fibers shed from electric clothes dryers (Xu et al., 2020). Microplastic particles have been detected in wide range of environments such as marine water, freshwater (Rodrigues et al., 2018), agroecosystems (Ng et al., 2018), terrestrial systems (Rillig, 2012), food and drinking water (Almaiman et al., 2021; Smith et al., 2018).

Although MPs have been studied more in aquatic environments, there is a growing acknowledgement that they are also present in terrestrial environments (Wright et al., 2020). MPs are deposited in the soil from water and wind originating from different sources including consumer products, the deterioration of plastic in landfills, wastewater sludge deposited on agricultural soils (Corradini et al., 2019), or car tire wear (Jan Kole et al., 2017). A review by Koutnik et al. (2021) analyzed the concentrations of MPs across 117 studies and showed that the concentration of MPs in soil varies by up to 8 orders of magnitude between locations tested with concentrations of ∼13,000 MP items/kg of dry soil found in agricultural and horticultural sites. In agricultural and natural soils, the most common polymer types of MPs are polyethylene (PE), polystyrene (PS), polypropylene (PP), polyvinyl chloride (PVC), and polyethylene terephthalate (PET) (Xu et al., 2020). PE, PP, and PET were found in nearly 80% of studies and in highest concentrations closer to urban areas. Soil and sediment samples from areas closer to industrial emission sites were found to have even higher concentrations of MPs with as many as 690,000 MP items/kg found in soil near industrial areas in China (Büks & Kaupenjohann, 2020). While these studies demonstrate the range of MPs in terrestrial environments, there are very few studies that examine how these levels may affect biota.

While MPs are abundant in terrestrial environments, most studies to date have focused on MP effects on aquatic animals (e.g., Cole & Galloway, 2015; Gaspar et al., 2018; Li et al., 2020; Lindeque et al., 2020; Mathalon & Hill, 2014; Sussarellu et al., 2016). MPs are readily ingested by a variety of aquatic animals (e.g., invertebrates, sea birds, fish) and have a wide range of direct and indirect toxicities (Anbumani & Kakkar, 2018; Huang et al., 2021; Lusher et al., 2017; McCauley & Bjorndal, 1999; Phillips & Bonner, 2015; Provencher et al., 2010; Rochman et al., 2014; Rummel et al., 2016; Ryan et al., 1988). Adverse effects of MPs on organisms can be categorized as either physical effects (caused by particle shape, size, or concentration), or chemical effects (caused by chemical additives originating from the plastic or absorbed from the surrounding environment). Chemical additives, such as BPA (bisphenol A), are added to plastic products during manufacturing to give plastic qualities such as color and transparency, or to improve its resistance to abiotic and biotic degradation (Hahladakis et al., 2018). For example, PVC commonly has phthalates added to improve flexibility and transparency. Phthalate molecules are not chemically bound to PVC, but physically interact through van der Waals forces, so they can easily diffuse into their surrounding environment through the air, water, or soil (Henkel et al., 2019; Zhang & Chen, 2014). Since MPs have hydrophobic properties, their surface can also adsorb and concentrate hydrophobic organic pollutants from the environment, such as organochlorine pesticides or polycyclic aromatic hydrocarbons (Zhang & Chen, 2014). Further, MPs can adsorb heavy metals such as cadmium, zinc, nickel, or lead from their immediate environment, or lead to the formation of biofilms (Kirstein et al., 2016) which may contribute a further toxicity of their own (Brennecke et al., 2016). Therefore, it is important to consider MPs a complex suite of contaminants (Rochman et al. 2019). To make matters more complicated, MPs that organisms can encounter vary in a wide range of shapes (e.g. fragments, foams, films, pellets and most commonly fibres), sizes, polymer types, and concentrations (Campanale et al., 2020).

We currently lack a full understanding of whether/how much microplastic ingestion impacts terrestrial animals, but research to date has demonstrated varied adverse effects of MPs on digestion, growth, reproduction, and behaviour. When the terrestrial snail *Achatina fulica* were fed PET microplastic fibres, the snails consumed less food and the fibres caused significant damage to the gastrointestinal tract (Song et al., 2019). By contrast, giant snails (*Achatina reticulata*) ingesting irregularly shaped and sized PET-MPs at 1 and 10% w/w grew larger and more quickly (de Felice et al., 2021). Springtails (*Folsomia candida*) exposed to 1% w/w PE-MPs in the soil suffered a 70.2% reduction in offspring produced (Ju et al., 2019). Microplastic ingestion similarly caused damage to male earthworm (*Eisenia andrei*) reproductive organs (inhibiting spermatogenesis) but caused negligible damage to female reproductive tissues, indicating that MP effects may be sex-specific (Kwak and An, 2021). Importantly, these studies specifically observed effects of untreated MP particles on terrestrial animals. Understanding the toxicological effects of untreated MPs alone is a prerequisite to a clear understanding of the additive or synergistic effects that might result from additional toxins bound to MPs.

Diet quality and composition affects the growth of insects (Nijhout & Callier, 2015), so the presence of MP in the diet may influence their growth rate and final size. To determine the physiological mechanisms underlying plastic effects on growth, we must first characterize those effects. Plastics may cause blockages in the gut of insects, leading to a decrease in the capacity for nutrient intake, and thus a reduction in adult body size. If insects can receive adequate nutrition while MPs are in their diet, they may also experience a toxicological effect on their growth or metabolic pathways (e.g., through endocrine disruption) due to harmful chemicals leaching out of the plastic. Any of these potential effects are expected to slow growth, reduce final body size, and (if severe enough) lead to mortality.

We hypothesized that as in earthworms, MP consumption is detrimental to insect growth and development. To test this, we used the tropical house cricket (*Gryllodes sigillatus*) as a model species. Cricket species have been used extensively as a model in previous studies to analyze dietary effects on growth and allometry (Bertram et al., 2021; Kelley & L’Heureux, 2021; Estenia & Gray, 2011). The effects of microplastic ingestion on growth may affect certain organs, such that changes in organ growth are more pronounced than changes in total body size (Whitman, 2008). As a larger insect with allometric growth, *G. sigillatus* are a prime candidate for studying changes in growth as a consequence of microplastic ingestion.

Here, we exposed crickets to either fluorescent PE microplastic beads (2.5, 5, or 10% w/w) or PET microfibers (0.25, 0.5 or 1% w/w) mixed into feed. PE and PET were chosen as they are two of the largest components of MP waste (Hamid et al., 2018). Ingestion of the MPs was confirmed by visual inspection of frass, while growth rate was measured through weekly mass and body length measurements. If the microplastics are being ingested by the crickets, we expected that increasing amounts of MP would lead to greater mortality or reduce the rate of body mass and/or size increase over developmental time.

## Methods

### Microplastics

Two types of MPs were tested in this experiment. Fluorescent PE microspheres (item #: UVPMS-BB-1.13 90-106um - 10g, Cospheric LLC, Santa Barbara, USA; 90-106μm, 1.10-1.14g/cc mean density, peak emission 445 nm when excited at 407 nm) were chosen as PE is one of the most used polymers in plastic material production (Horton et al., 2017), and the fluorescence allowed relatively simple tracking of bead ingestion during the experiment. The second type of plastic used was untreated spun PET (item #777, spun polyester type 54; Testfabrics, Inc., West Pittston, PA, USA). PE and PET are both used commonly in aquatic experiments, allowing direct comparisons to previous studies. The PET fabric was cut into small pieces with fabric scissors and blended using a Magic Bullet (nutribullet, LLC, Pacoima, California, USA) to simulate microfibers produced by domestic dryers and fabric industries (Kapp & Miller, 2020). The fabric was blended in three 5-6 second intervals and stored in an air-tight container until use to prevent air-bourne contamination from other sources of MP.

### Cricket rearing

*Gryllodes sigillatus* eggs were supplied by a commercial insect farm (Entomo Farms; Norwood, Canada) and placed directly into an incubator maintained at 32 ± 2°C and 35 ± 5% RH on a 14:10 L:D cycle. Eggs were moistened with water and gently stirred every other day to prevent desiccation and mold growth until emergence. Once emerged, individual crickets were housed in 3 ¼ oz plastic solo cups with matching plastic lids and provided a piece of egg carton for shelter along with food and water *ad libitum* for the duration of the experiment. Because each cricket was reared under the same conditions, any effect of plastic contamination from their housing would be the same among treatment groups. Regardless, plastic solo cups used for cricket housing were washed and dried between feeding to prevent further plastic contamination. Water for crickets was provided in 0.65 mL microcentrifuge tubes with moistened dental cotton covering the opening and the base diet consisted of a proprietary mixture of corn, soybean, herring, and hog meal. Food and water were replaced every 3-4 days for each cricket for the duration of the experiments.

### Microplastic Bead Feeding

Within 24 h of emergence, 96 newly emerged cricket nymphs were selected at random and placed into individual housing. Crickets were divided into four treatment groups with 24 crickets in each group. Each group received either 0 (control), 2.5, 5, or 10% w/w fluorescent blue microplastic beads mixed into the dry feed (Figure 1).

**Figure 1:**
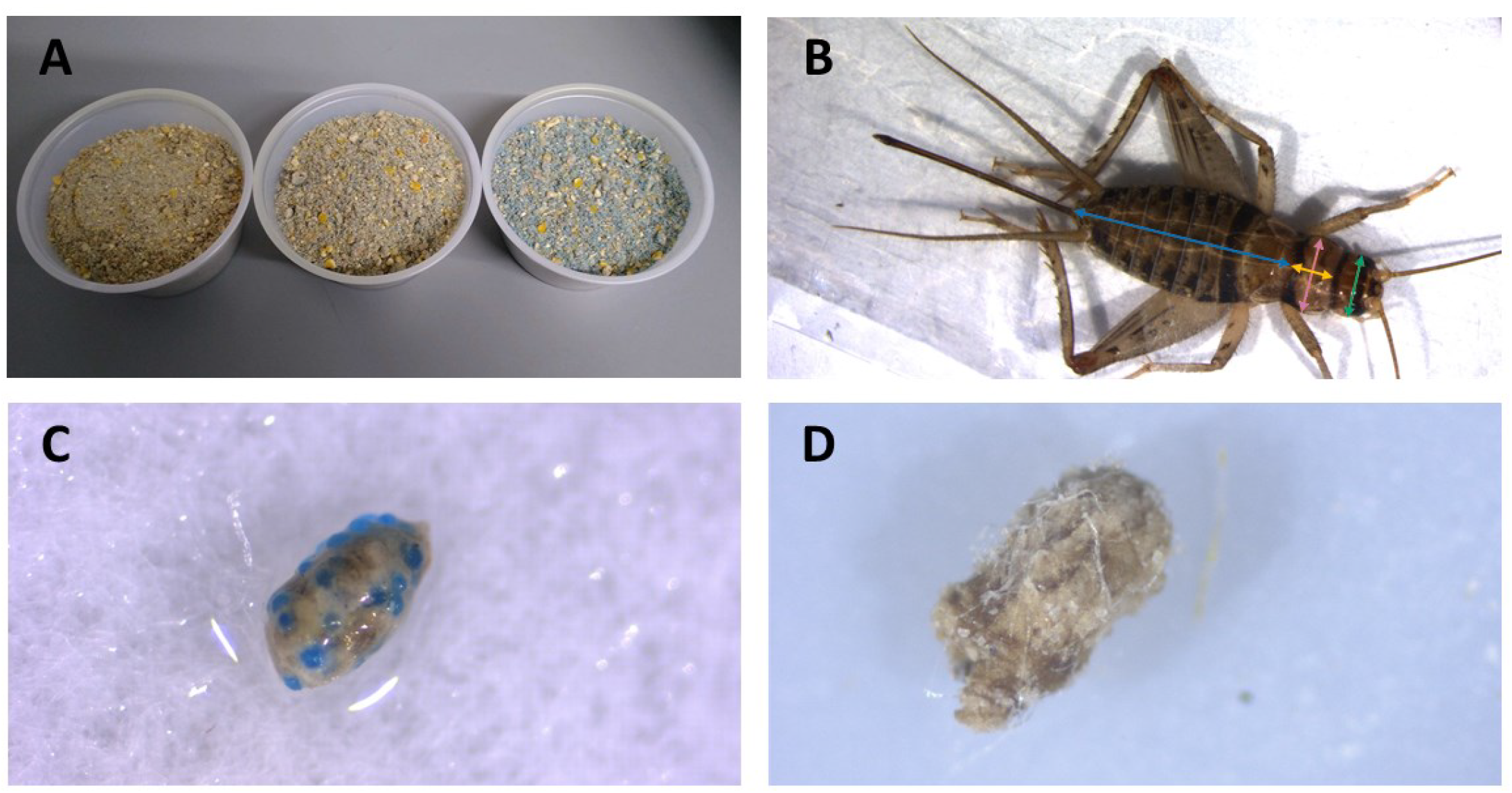
Cricket diet mixed with blue, fluorescent microplastic beads. Concentrations shown from left to right are 2.5, 5 and 10% w/w plastic mixed with base diet (A). Example of measurements taken from crickets. Blue = abdomen length, yellow = thorax width, pink = thorax length, green = head width (B). Frass from crickets feeding on 10% w/w fluorescent polyethylene microbeads in feed (C) and frass from crickets feeding on 1% w/w polyethylene terephthalate in feed (D).

### Microplastic Fiber Feeding

A second experiment was conducted with PET microfibers. Within 24 h of emergence, 96 newly emerged cricket nymphs were selected at random and placed into individual housing. Crickets were divided into four treatment groups with 24 crickets in each group. Each group received either 0 (control), 0.25, 0.5, or 1% w/w PET microfibers mixed into the wet feed. A lowered concentration of microfibers was used to adjust for the difference in density of plastics. PET fibers were dispersed in water before mixed into feed to ensure an even distribution of fibers.

### Cricket Body Measurement

Body mass and size of crickets were measured weekly starting 24 h after cricket nymph emergence. Body mass was measured by placing live crickets into a pre-weighed 2 Ml microcentrifuge tube and weighing them with a Sartorius ME-5 microbalance scale (Precision Weighing Balances, Bradford, MA, USA). To measure body size, live crickets were placed into clear, flat 2.5” x 3.5” plastic bags, and photographed using a pre-calibrated Stemi 508 trinocular dissecting microscope equipped with a camera (Zeiss, Jena, Germany). A scale bar was photographed with each cricket to accurately perform digital measurements. Digital measurements were then taken from the photos using ImageJ v.148 (National Institutes of Health, Bethesda, MD, U.S.A). Head width (maximal distance between the outer edges of the eyes), pronotum width (maximal distance across the coronal width of the pronotum) and length (maximal distance down the sagittal length of the pronotum), and abdomen length (maximal distance down the sagittal length of the abdomen) (Figure 1) were measured for each individual over the duration of eight weeks. Sex of the crickets was determined at the 5-6 weeks post emergence, when the ovipositor was present in female crickets. Male crickets lack an ovipositor and have pronounced wings on their abdomen that are used to produce their characteristic mating calls. Crickets were euthanized by freezing after the final measurements.

### Data Analysis

Data analysis was conducted in R version 4.0.2 using R Studio version 1.3.1073 (R Core Team, 2020). Data distributions and variance were assessed using Shapiro-Wilk tests and Q-Q plots. To increase confidence in our findings and test for interactive effects of developmental time and MP concentration on body size and mass, the effects of MP fiber and bead concentration in feed on the growth of the crickets were analyzed using linear mixed effects models with the lme() function in R (Bates et al. 2014). In addition, to test specifically for an effect of plastic feeding on final body mass and size, the final week of measurements were analyzed using a Kruskal-Wallis test with the kruskal.test() function in R as assumptions of data normality could not be met with data transformation. Cricket measurements were separated by sex to account for any differences in size caused by sexual dimorphism in *G. sigillatus* (Archer et al., 2012). The week of each measurement was treated as a fixed effect while each individual cricket measured was treated as a random effect to account for variability in growth per individual.

## Results

### Microbead Ingestion

40 male and 39 female crickets total from all treatment groups survived for the duration of the experiment while consuming different concentrations of fluorescent PE microplastic beads in their diet (Figure 2 and 3). The majority of crickets that did not survive died within the first week of growth (Table 1). Subsequent loss of crickets during the study were caused by crickets escaping or accidental damage caused during handling (one animal fed the 5% w/w diet and one control). The number of crickets that died during the experiment were not statistically significantly different from the control. Contrary to our predictions, the growth of both males and females did not significantly change in any of the parameters we tested (Supplementary Tables 1 and 2). Likewise, the growth of males and females was not significantly different at any concentration of PE-MP beads in feed when tested with a Kruskal-Wallis test.

**Figure 2:**
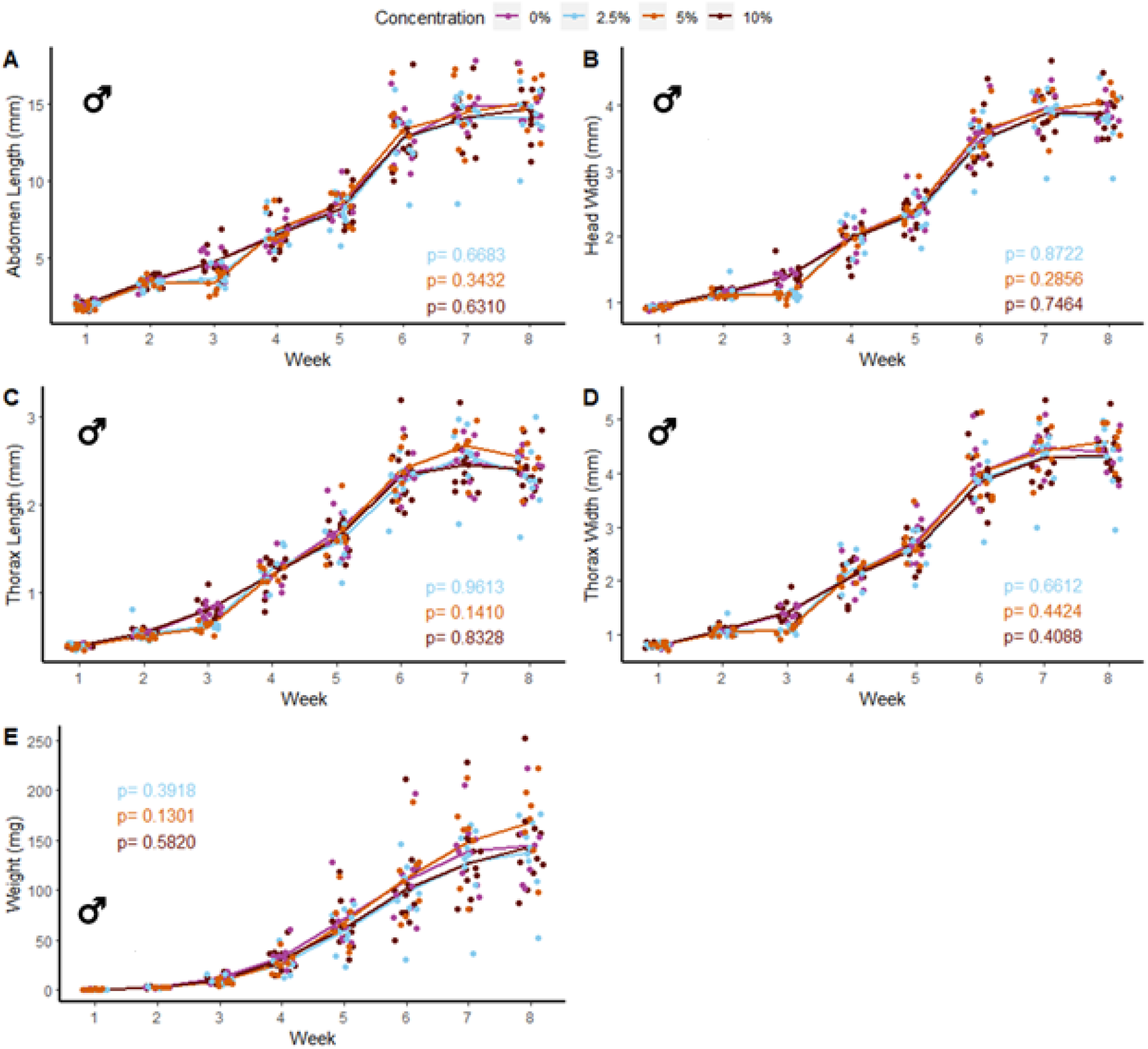
Body size and mass of male *Gryllodes sigillatus* fed a diet containing PE microbeads. Initial measurements of crickets were taken once cricket nymphs emerged from eggs at week 1. Points represent individual cricket measurements and lines represent mean values between measurements. P-values are presented as an interaction of the concentration of PE microbeads with the week of growth. Each letter indicates a different type of measurement with A = abdomen length, B = head width, C = thorax length, D = thorax width, E = weight.

**Figure 3:**
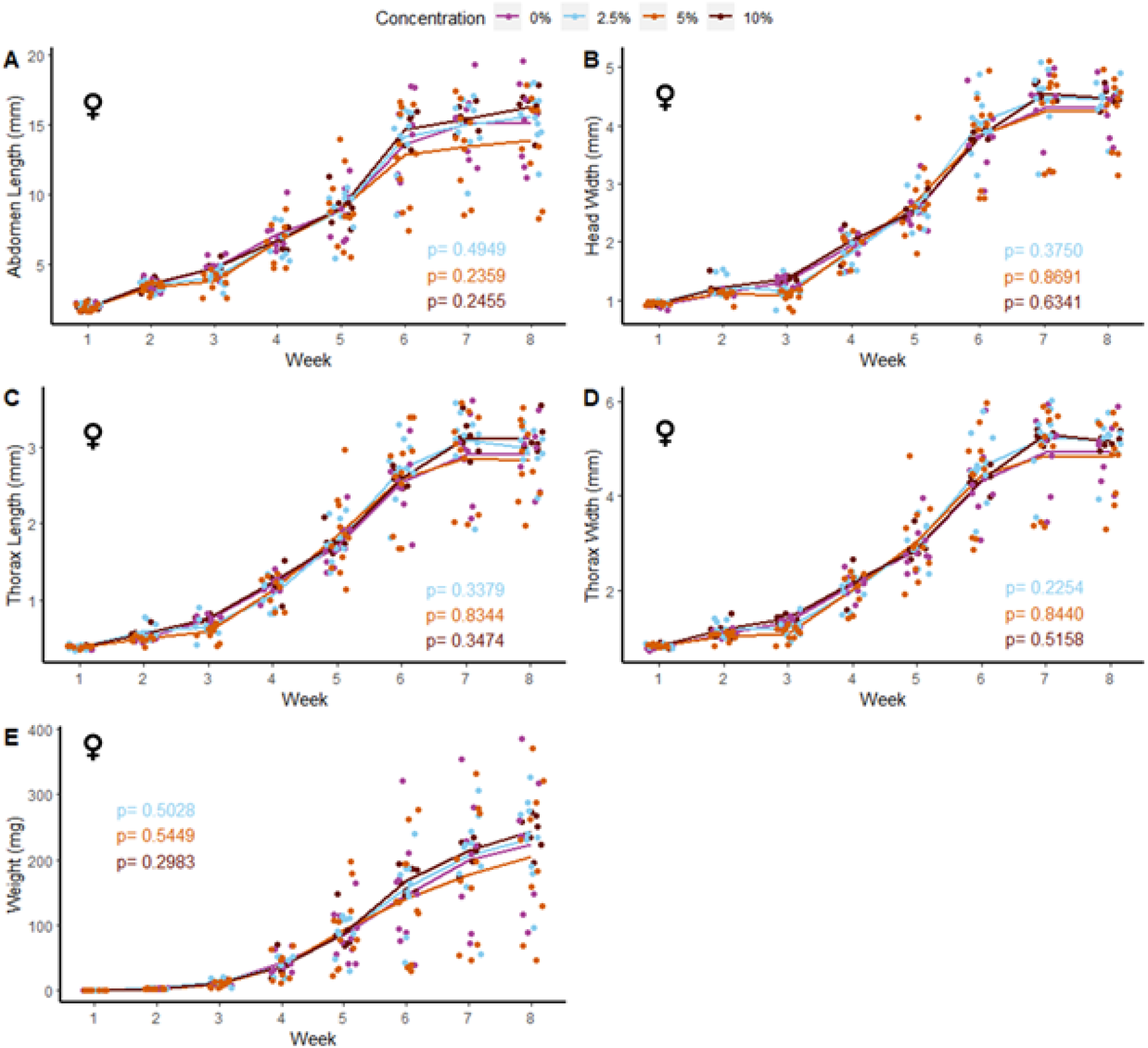
Body size and mass of female *Gryllodes sigillatus* fed a diet containing PE microbeads. Initial measurements of crickets were taken once cricket nymphs emerged from eggs at week 1. Points represent individual cricket measurements and lines represent mean values between measurements. P-values are presented as an interaction of the concentration of PE microbeads with the week of growth. Each letter indicates a different type of measurement with A = abdomen length, B = head width, C = thorax length, D = thorax width, E = weight.

**Table 1:** Mortality of *Gryllodes sigillatus* during MP feeding experiments. Crickets that died or were lost during measurement were excluded from the experiment. N (Start): Number of crickets at the beginning of the experiment. N (Died): Number of crickets that died during the experiment. Mortality: Proportion of crickets that died during the experiment.

| Microbead<br>concentration (% w/w) | N (Start) | N (died) | Mortality | Average age at<br>death (weeks) |
| --- | --- | --- | --- | --- |
| **Microbeads** |  |  |  |  |
| 0 | 24 | 5 | 0.21 | 2 |
| 2.5 | 24 | 5 | 0.21 | 2 |
| 5 | 24 | 6 | 0.25 | 2 |
| 10 | 24 | 4 | 0.17 | 2 |
| **Microfibers** |  |  |  |  |
| 0 | 24 | 1 | 0.04 | 2 |
| 0.25 | 24 | 0 | 0.00 | 2 |
| 0.5 | 24 | 5 | 0.21 | 2 |
| 1 | 24 | 4 | 0.17 | 2 |

### Microfiber Ingestion

47 male and 37 female crickets total from all treatment groups survived for the duration of the experiment while consuming different concentrations of PET microfibers in their diet (Figure 4 and 5). The majority of crickets that did not survive died within the first or second week of growth. Unlike the microbead mortality, crickets fed 0.5 and 1% microfiber diets (the two highest exposure levels) tended to die more frequently than those fed the other diets, but too few animals died during the experiment to reliably test this result using a statistical analysis (Table 1). Subsequent loss of crickets during the study were caused by crickets escaping or damage caused during handling (three animals fed 1% w/w diet, one animal fed 0.5% w/w diet). There were no significant interactive effects of age and plastic dose on any measurement in males with the exception of 0.5% w/w PET on body mass. Males fed 0.5% w/w PET showed a significant decrease in mass (LME; *t*(321) = - 2.4, *p* = 0.0301) (Supplementary Table 3). Female abdomen length showed the largest reduction in mean size for females fed 1% PET compared to the control (LME; *t*(245) = -4.344, *p* = < 0.0001) (Supplementary Table 4). Similarly, thorax width was significantly reduced in female crickets fed a 1% w/w PET microfiber diet (LME; *t*(245) = -3.4, *p* = 0.0009). The 1% w/w PET microfibers diet also significantly reduced the thorax length (LME; *t*(245) = - 0.05, *p* = 0.0041) and head width (LME; *t*(245) = - 0.04, *p* = 0.0172). Overall, higher microfiber doses significantly reduced body mass in female crickets (LME; *t*(245) = -12.3, *p* = 0.0002).

**Figure 4:**
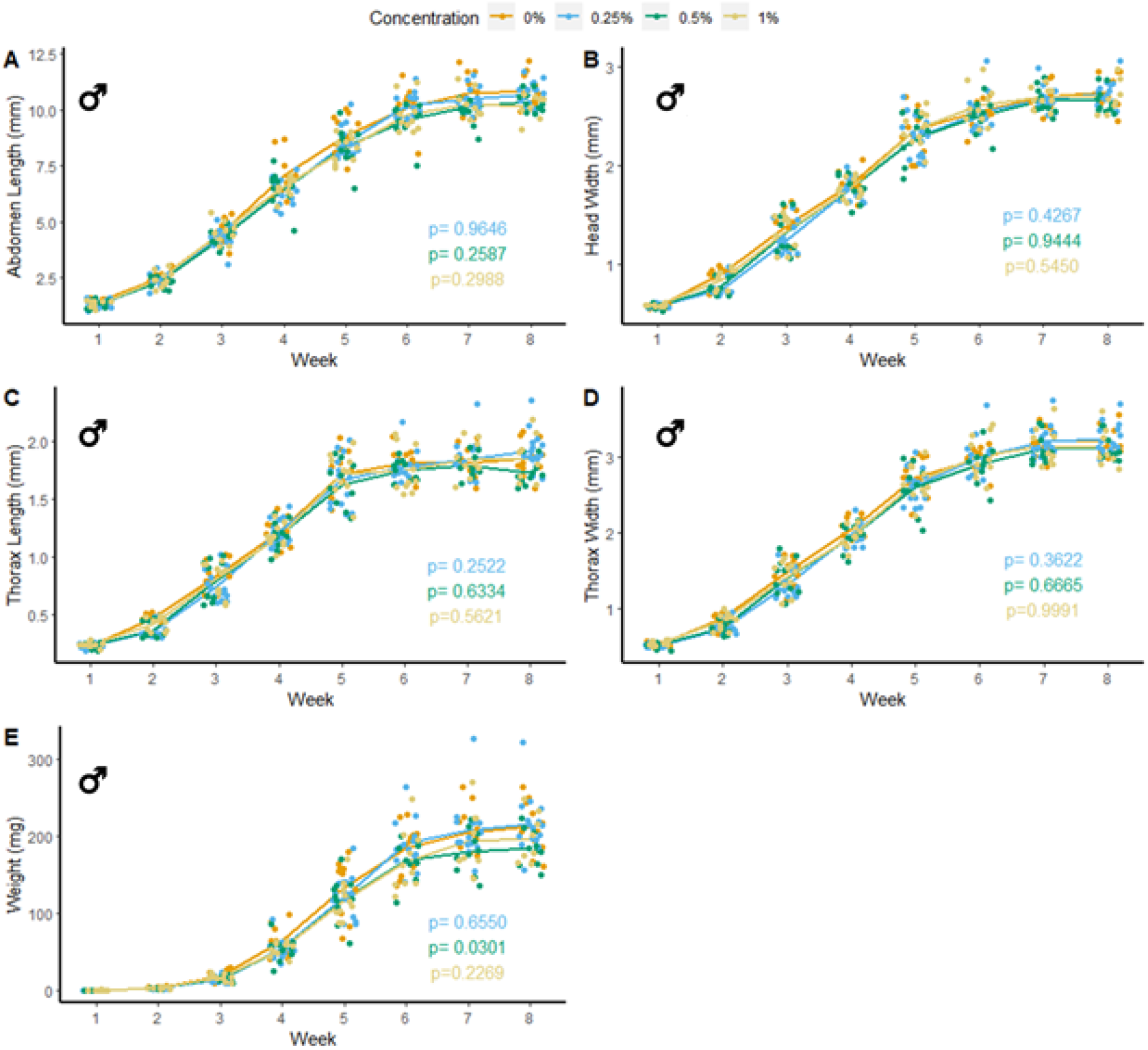
Body size and mass of male *Gryllodes sigillatus* fed a diet containing PET microfibers. Initial measurements of crickets were taken once cricket nymphs emerged from eggs at week 1. Points represent individual cricket measurement and lines represent mean values between measurements. P-values are presented as an interaction of the concentration of PET microfiber with the week of growth. Each letter indicates a different type of measurement with A = abdomen length, B = head width, C = thorax length, D = thorax width, E = weight.

**Figure 5:**
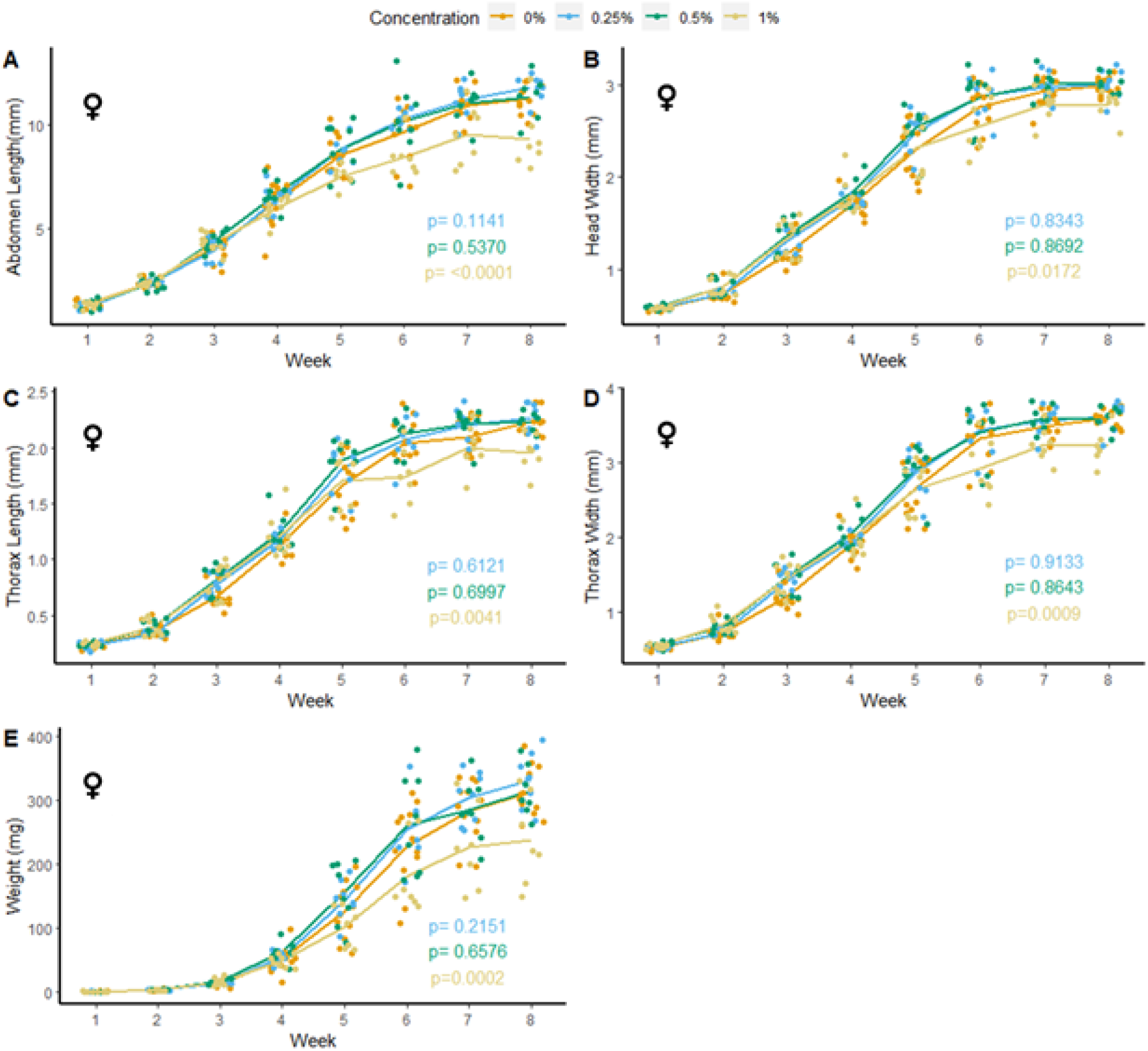
Body size and mass of female *Gryllodes sigillatus* fed a diet containing PET microfibers. Initial measurements of crickets were taken once cricket nymphs emerged from eggs at week 1. Points represent individual cricket measurement and lines represent mean values between measurements. P-values are presented as an interaction of the concentration of PET microfiber with the week of growth. Each letter indicates a different type of measurement with A = abdomen length, B = head width, C = thorax length, D = thorax width, E = weight.

Following the results of the linear mixed effects model, a Kruskal-Wallis test was performed to validate the significance seen from the non-normal results. Based on this analysis, female crickets fed 1% w/w PET microfibers were significantly different from the control at their final week of growth measurement. High concentrations of PET microfibers caused a significant decrease in abdomen length (KW: χ2 = 10.4, *p* = 0.0152). Thorax length was significantly reduced by the 1% w/w PET microfiber diet (KW: χ2 = 11.9, *p* = 0.0078). As well, consumption of high concentrations of PET microfibers significantly reduced thorax width (KW: χ2 = 10.4, *p* = 0.0155) and head width was significantly smaller when feeding on the 1% w/w PET microfiber diet (KW: χ2 = 12.0, *p* = 0.0075). Lastly, female crickets weighed significantly less than the control when fed 1% w/w PET microfibers (KW: χ2 = 7.7 *p* = 0.0522). Taken together, these KW test results support the outcomes of our LME-based approach, and verified that abdomen length, thorax length, thorax width, and mass of female crickets fed 1% w/w PET microfibers differ significantly from the control at the end of their developmental period.

## Discussion

MP contamination now permeates nearly every environment in the world and has become a major environmental concern (Kumar et al., 2021). We present a novel study that investigates the effects of microplastic ingestion on the growth and development of a hemimetabolous terrestrial insect. As with previous MP studies on other model animals, the crickets used in our experiment were found to ingest and pass the plastics presented in their food via their gastrointestinal tract (Immerschitt & Martens, 2020; Windsor et al., 2019; Wright et al., 2013). We found that when both male and female *G. sigillatus* ate high concentrations of PE beads, there was no change in body size, weight, or mortality. By contrast, when female crickets were fed high concentrations (1% w/w) of PET microfibers, they experienced a significant reduction in total body size and weight throughout development, while males only experienced a reduction in weight at 0.5% w/w PET, but not any measure of body size.

Further questions are raised regarding the mechanisms through which MPs reduce the growth of crickets and the toxicological effects caused by PET or PE. Because the PE microbeads did not result in any changes in body size, it is possible the decrease in body size due to PET microfibres is caused by the morphology of the plastic (fiber vs. bead). Goldfish (*Carassius auratus*) chewed and often rejected MP fragments and pellets, but passively ingested MP fibers, which caused significant damage to the gastrointestinal tract and the liver of the goldfish (Jabeen et al., 2018). PE-MPs did not cause significant effects on the growth, survival, or reproduction of the worm, *Eisenia andrei*, but did show evidence of physical damage to the gut (Rodriguez-Seijo et al., 2017). Because *G. sigillatus* readily ingested the beads, they may be causing limited physical damage to the gut walls, however, beads were excreted and passed completely through the digestive system.

We were surprised that crickets ingesting the MP beads could maintain similar growth rates when their diet was diluted with up to 10% indigestible plastic. We assumed that a high concentration of MP beads would result in a low-quality diet and thereby impact growth. In the grasshopper, *Melanopus differentialis*, a lower quality diet led to greater gut size (Yang & Joern, 1994). Changes in gut size may allow for insects to ingest more food at one point in development or increase the efficiency of digestion from a longer retention of food in the gut. As such, *G. sigillatus* in the present study may have compensated for the plastic bead ingestion by adjusting the size of their gut and thereby allowing more food to be retained and digested per unit time. By contrast, microfibre feeding did decrease female growth rates. This difference in the impacts of beads and fibres supports the notion that plastic form is critical to understanding its impacts on animals. We did not collect data on rates of food ingestion, but suspect that the microfibers may be causing a blockage in the cricket gut. Food was provided *ad libitum* for the crickets; therefore, they may have compensated for the lack of nutrition through increased feeding. *G. sigillatus* may compensate for the diluted nutrients in their food by increasing their food intake to maintain key biological functions.

*G. sigillatus* ate PE microbeads indiscriminately when they were presented in their food around their third week of age. MPs in the environment are found in a wide range of sizes and *G. sigillatus* may avoid MP in their diet if presented in these different sizes (Allison, unpub data). Likewise, if provided an alternative food source simultaneously without the presence of MPs, *G. sigillatus* may also show a preference towards a diet free of plastic. The freshwater shrimp, *Gammarus pulex*, avoided a diet with acrylic microfibers when presented with an alternative diet without microfibers (Yardy & Callaghan, 2020). In our experiments, *G. sigillatus* were given no choice but to consume a diet containing plastics, and since they readily ingested diets with MPs, crickets proved to be an ideal model system to examine the effects of plastic size, shape, and polymer type on insect feeding preferences. If given a choice, *G. sigillatus* may choose to avoid food contaminated with MPs or they may be unable to detect MPs present in their diet.

We observed no differences in the size of crickets fed PET-MP, but did detect a significant change in body weight in those fed the 0.5% w/w PET diet. This result may not be biologically relevant as it was only seen in one group of mass measurement, although changes in the fat stores may still be occurring in the cricket population to maintain their growth trajectory. The insect fat body is important to maintain metabolic homeostasis. It functions as storage for excess nutrients and synthesizes lipids, glycogen, and most of the hemolymph proteins and circulating metabolites (Arrese & Soulages, 2010). When insects experience periods of starvation, they undergo an increase in hemolymph lipid concentration due to an elevation in lipids or diglycerides produced from the stored fat body (Beenakkers et al., 1985). If *G. sigillatus* are experiencing a decreased nutrient intake due to high concentrations of PET providing no nutritional substance in their diet, then the concentration of lipids in the hemolymph and/or whole-body lipid stores could be measured to verify if reduced nutrient intake is driving the observed reductions in cricket body mass.

Although females fed 1% PET microfibers experienced a decrease in each of the parameters measured, abdomen length showed the greatest reduction in size on average in the female crickets (Figure 5a). This can be indicative of the microfibers depriving the female *G. sigillatus* of the energetic costs for the production of oocytes, resulting in the reduction in size (Ziegler & van Antwerpen, 2006). In another cricket species, *Gryllus bimaculatus*, fat body mass peaked after adult eclosion then depleted over 48 h following a 16-fold increase in ovary weight (Lorenz & Anand, 2004). The reduced weight seen in female *G. sigillatus* fed 1% microfibers may be an indicator of an insufficient production of fat body during development, which can later decrease the production of oocytes and thereby impact reproductive fitness (Ziegler & van Antwerpen, 2006). Abdomen size has also been directly correlated to reproductive output (Wickman & Karlsson, 1989), suggesting that smaller abdomen size in females can indicate a reduction in number of offspring. Future studies can investigate how production of eggs or viability of offspring are impacted by the same levels of MP ingestion used here. By measuring the number of eggs present in adult female crickets fed high concentrations of PET-MP fibers, one could see if the reduction in abdomen size translates to a reduction in reproductive output. In support of this hypothesis, female *Drosophila melanogaster* were recently shown to suffer a 50% decrease in egg oviposition when fed diets containing a concentration of 20 g/L PET-MP (Shen et al., 2021). reproduction was similarly inhibited in springtails (*Folsomia candida*) in the presence of increasing concentrations of PE-MPs in the soil (Ju et al., 2019). Although PE-MP ingestion did not lead to any significant decrease in the cricket’s abdomen length in the present study, ingestion of PE microbeads may or may not affect the reproductive output of crickets without influencing body size. *G. sigillatus* undergo multiple matings and lay multiple batches of eggs throughout adulthood (Sakaluk et al., 2002). Since *G. sigillatus* readily consume MPs, like *Drosophila* (Shen et al., 2021) they could provide an ideal study system for the reproductive or multi-generational effects that MP ingestion may have on insect fitness.

This experiment specifically isolated the effects of untreated PE and PET microplastics on the growth of the cricket *G. sigillatus*. The effects of both the microbead and microfiber feeding observed here represent a first step to understanding how microplastics affect terrestrial species in isolation. As such, it may not be completely indicative of how MPs ingested outside of a laboratory setting can affect *G. sigillatus* or other terrestrial species. The toxicity of plastics can be significantly altered though contact with other organic pollutants that can bind both MP particles and fibers (Vázquez & Rahman, 2021). For example, the most common type of litter on Earth, cigarette butts, potentially release about 0.3 million cellulose acetate microfibers a year with toxic leachates absorbed from the cigarette smoke (Belzagui et al., 2021). Because of MPs’ readiness to leach different contaminants from the environment in addition to the number of additives used in the production of plastic, it is unlikely that these animals will experience exposure to virgin, uncontaminated MPs in the wild (Hahladakis et al., 2018), and yet clearly these plastics alone have significant effects on growth. Zhang et al. (2020) investigated damage caused to fruit flies (*Drosophila melanogaster*) by not only MPs, but a combination of MPs and cadmium (Cd), a widely used heavy metal pigment that readily leaches out of plastic. The combination of Cd mixed with PS microbeads intensified the toxicity of the heavy metal, shown through reduced climbing activity and increased damage to their gut. Since Cd is only one of the toxins that can be readily leached from MPs, characterizing and understanding the toxicity of MP remains a huge challenge. Regardless, virgin MPs can have their own toxicological effects on biota that we must first understand before we can adequately comprehend such the physiological causes and consequences of any interactive effects observed.

Our results revealed that high (1% w/w) concentrations of untreated PET microfibers can significantly hinder the growth of female *G. sigillatus*, and possibly the weight, but not size, of males. The physiological mechanisms which cause this change in growth and how these mechanisms might interact with other toxins bound to MPs remain to be determined, but we argue that crickets are an excellent study system in which to explore them. To our surprise, PE microbead ingestion caused no detectable change in the growth of both male and female crickets, which suggests that MP form is important to consider in laboratory experiments of plastic effects, and that insects may compensate for reduced nutrient content in the diet by ingesting more food. Questions remain about how plastics can be transformed in the gut of terrestrial insects and whether there are other effects of MP feeding on insect fitness that cannot be observed through body size or weight measurements, such as effects on reproduction or behaviour.

## Author Contributions

All authors together conceived the study and designed the experiments. S.F. and M.R. carried out the experiments. S.F. ran the systematic literature search and developed all tables. S.F. curated and analyzed the data and created visualizations. H.A.M. provided resources and supervision. S.F. drafted the manuscript and all authors edited the manuscript.

## Acknowledgments

I would like to thank my committee members Kyle Biggar and Kathleen Gilmour for their support and guidance during my research. As well, thank you to Izzy Munevar and Emily McCoville for help with rearing crickets and measurements. Thank you to Lisa Erdle for her support provided on the handling and processing of plastic microfibers for use in insect ingestion experiments.

## Data Availability Statement

All data is provided as a supplementary file for review and the same file will be included as supplementary material should the manuscript be accepted for publication.

