## Supplementary materials for "Ingestion of microplastic fibres, but not microplastic beads, impacts growth rates in the tropical house cricket *Gryllodes sigillatus*"

**Table S1:** Results of linear mixed effects model for microbead concentration vs. growth of the abdomen, thorax, head, and weight of female *G. sigillatus* with cricket identity as random effects and presented with interactions with the cricket age (week). Results considered statistically significant from the control ($p < 0.05$) are presented in bold.

| | Female Microbead Diet | | | | | | | | |
|---|---|---|---|---|---|---|---|---|---|
| | Weight | | | Head Width | | | Thorax Width | | |
| *Coeffcient* | *Estimates* | *Conf. Int (95%)* | *P-Value* | *Estimates* | *Conf. Int (95%)* | *P-Value* | *Estimates* | *Conf. Int (95%)* | *P-Value* |
| (Intercept) | -69.62 | -98.72 – -40.52 | **<0.001** | -0.01 | -0.27 – 0.25 | 0.947 | -0.3 | -0.64 – 0.03 | 0.074 |
| Concentration [2.5%] | -3.5 | -45.69 – 38.68 | 0.867 | -0.05 | -0.43 – 0.32 | 0.772 | -0.1 | -0.58 – 0.38 | 0.673 |
| Concentration [5%] | 5.57 | -35.75 – 46.90 | 0.786 | -0.04 | -0.41 – 0.33 | 0.84 | -0.05 | -0.52 – 0.43 | 0.845 |
| Concentration [10%] | -4.49 | -49.88 – 40.90 | 0.842 | 0.03 | -0.38 – 0.44 | 0.872 | 0.01 | -0.51 – 0.53 | 0.964 |
| week | 34.92 | 30.40 – 39.45 | **<0.001** | 0.56 | 0.52 – 0.61 | **<0.001** | 0.69 | 0.63 – 0.75 | **<0.001** |
| Concentration [2.5%] * week | 2.11 | -4.09 – 8.31 | 0.503 | 0.03 | -0.04 – 0.10 | 0.375 | 0.05 | -0.03 – 0.13 | 0.225 |
| Concentration [5%] * week | -1.89 | -8.03 – 4.25 | 0.545 | 0.01 | -0.06 – 0.07 | 0.869 | 0.01 | -0.07 – 0.09 | 0.844 |
| Concentration [10%] * week | 3.57 | -3.17 – 10.31 | 0.298 | 0.02 | -0.06 – 0.09 | 0.634 | 0.03 | -0.06 – 0.12 | 0.516 |
| **Random Effects** | | | | | | | | | |
| σ2 | | 1938.81 | | | 0.23 | | | 0.35 | |
| τ00 | | 944.30 $_{\text{cricket}}$ | | | 0.03 $_{\text{cricket}}$ | | | 0.06 $_{\text{cricket}}$ | |
| ICC | | | | | 0.12 | | | 0.16 | |
| N | | 39 $_{\text{cricket}}$ | | | 39 $_{\text{cricket}}$ | | | 39 $_{\text{cricket}}$ | |
| Observations | | 289 | | | 289 | | | 289 | |
| Marginal $R^2$ / Conditional $R^2$ | | 0.780 / NA | | | 0.874 / 0.889 | | | 0.868 / 0.888 | |

| | Female Microbead Diet | | | | | |
|---|---|---|---|---|---|---|
| | Thorax Length | | | Abdomen Length | | |
| *Coeffcient* | *Estimates* | *Conf. Int (95%)* | *P-Value* | *Estimates* | *Conf. Int (95%)* | *P-Value* |
| (Intercept) | -0.28 | -0.48 – -0.08 | **0.006** | -0.81 | -1.89 – 0.27 | 0.142 |
| Concentration [2.5%] | -0.05 | -0.34 – 0.24 | 0.752 | -0.34 | -1.91 – 1.22 | 0.659 |
| Concentration [5%] | -0.03 | -0.31 – 0.26 | 0.853 | 0.09 | -1.45 – 1.62 | 0.911 |
| Concentration [10%] | -0.01 | -0.33 – 0.30 | 0.93 | -0.34 | -2.02 – 1.35 | 0.685 |
| week | 0.42 | 0.38 – 0.46 | **<0.001** | 2.12 | 1.94 – 2.30 | **<0.001** |
| Concentration [2.5%] * week | 0.02 | -0.03 – 0.07 | 0.338 | 0.09 | -0.16 – 0.33 | 0.495 |
| Concentration [5%] * week | 0.01 | -0.04 – 0.05 | 0.834 | -0.15 | -0.39 – 0.10 | 0.236 |
| Concentration [10%] * week | 0.03 | -0.03 – 0.08 | 0.347 | 0.16 | -0.11 – 0.43 | 0.246 |
| **Random Effects** | | | | | | |
| σ2 | | 0.13 | | | 3.1 | |
| τ00 | | 0.02 $_{\text{cricket}}$ | | | 1.02 $_{\text{cricket}}$ | |
| ICC | | 0.16 | | | 0.25 | |
| N | | 39 $_{\text{cricket}}$ | | | 39 $_{\text{cricket}}$ | |
| Observations | | 289 | | | 289 | |
| Marginal $R^2$ / Conditional $R^2$ | | 0.872 / 0.892 | | | 0.856 / 0.892 | |

**Table S2:** Results of linear mixed effects model for microbead concentration vs. growth of the abdomen, thorax, head, and weight of male *G. sigillatus.* cricket identity as random effects and presented with interactions with the cricket age (week). Results considered statistically significant from the control ($p$ < 0.05) are presented in bold.

| | Male Microbead Diet | | | | | | | | | | | | | | |
|---|---|---|---|---|---|---|---|---|---|---|---|---|---|---|---|
| | Weight | | | Head Width | | | Thorax Width | | | Thorax Length | | | Abdomen Length | | |
| *Coeffcient* | *Estimates* | *Conf. Int (95%)* | *P-Value* | *Estimates* | *Conf. Int (95%)* | *P-Value* | *Estimates* | *Conf. Int (95%)* | *P-Value* | *Estimates* | *Conf. Int (95%)* | *P-Value* | *Estimates* | *Conf. Int (95%)* | *P-Value* |
| (Intercept) | -43.22 | -58.36 – -28.07 | **<0.001** | 0.2 | 0.02 – 0.38 | **0.032** | -0.07 | -0.30 – 0.16 | 0.53 | -0.07 | -0.22 – 0.08 | 0.343 | -0.7 | -1.52 – 0.12 | 0.093 |
| Concentration [2.5%] | 0.87 | -21.75 – 23.49 | 0.938 | -0.05 | -0.32 – 0.22 | 0.713 | -0.04 | -0.38 – 0.31 | 0.827 | -0.04 | -0.26 – 0.18 | 0.72 | -0.19 | -1.41 – 1.03 | 0.751 |
| Concentration [5%] | -8.54 | -32.02 – 14.93 | 0.465 | -0.12 | -0.40 – 0.16 | 0.382 | -0.12 | -0.48 – 0.23 | 0.486 | -0.13 | -0.35 – 0.10 | 0.268 | -0.48 | -1.74 – 0.79 | 0.449 |
| Concentration [10%] | 0.04 | -20.70 – 20.77 | 0.997 | 0.02 | -0.23 – 0.26 | 0.902 | 0.05 | -0.27 – 0.36 | 0.767 | 0.01 | -0.19 – 0.20 | 0.951 | 0.11 | -1.01 – 1.23 | 0.844 |
| week | 23.83 | 21.51 – 26.15 | **<0.001** | 0.49 | 0.46 – 0.53 | **<0.001** | 0.6 | 0.56 – 0.64 | **<0.001** | 0.34 | 0.32 – 0.37 | **<0.001** | 2.04 | 1.90 – 2.18 | **<0.001** |
| Concentration [2.5%] * week | -1.45 | -4.77 – 1.87 | 0.392 | 0 | -0.06 – 0.05 | 0.872 | -0.01 | -0.07 – 0.05 | 0.661 | 0 | -0.04 – 0.04 | 0.961 | -0.04 | -0.25 – 0.16 | 0.668 |
| Concentration [5%] * week | 2.7 | -0.80 – 6.20 | 0.13 | 0.03 | -0.02 – 0.08 | 0.286 | 0.03 | -0.04 – 0.09 | 0.442 | 0.03 | -0.01 – 0.07 | 0.141 | 0.1 | -0.11 – 0.32 | 0.343 |
| Concentration [10%] * week | -0.86 | -3.94 – 2.21 | 0.582 | -0.01 | -0.05 – 0.04 | 0.746 | -0.02 | -0.08 – 0.03 | 0.409 | 0 | -0.04 – 0.03 | 0.833 | -0.05 | -0.24 – 0.14 | 0.631 |
| **Random Effects** | | | | | | | | | | | | | | | |
| $\sigma^2$ | | 550.14 | | | 0.13 | | | 0.19 | | | 0.08 | | | 2.09 | |
| $\tau_{00}$ | | 249.36 $_{cricket}$ | | | 0.00 $_{cricket}$ | | | 0.02 $_{cricket}$ | | | 0.01 $_{cricket}$ | | | 0.42 $_{cricket}$ | |
| ICC | | | | | 0.03 | | | 0.1 | | | 0.06 | | | 0.17 | |
| N | | 40 $_{cricket}$ | | | 40 $_{cricket}$ | | | 40 $_{cricket}$ | | | 40 $_{cricket}$ | | | 40 $_{cricket}$ | |
| Observations | | 306 | | | 306 | | | 306 | | | 306 | | | 306 | |
| Marginal $R^2$ / Conditional $R^2$ | | 0.845 / NA | | | 0.906 / 0.909 | | | 0.900 / 0.910 | | | 0.885 / 0.892 | | | 0.898 / 0.915 | |

**Table S3:** Results of linear mixed effects model for microfiber concentration vs. growth of the abdomen, thorax, head, and weight of female *G. sigillatus* with cricket identity as random effects and presented with interactions with the cricket age (week). Results considered statistically significant from the control ($p < 0.05$) are presented in bold.

| | Female Microfiber Diet | | | | | | | | | | | | | | |
|---|---|---|---|---|---|---|---|---|---|---|---|---|---|---|---|
| | **Weight** | | | **Head Width** | | | **Thorax Width** | | | **Thorax Length** | | | **Abdomen Length** | | |
| *Coeffcient* | *Estimates* | *Conf. Int (95%)* | *P-Value* | *Estimates* | *Conf. Int (95%)* | *P-Value* | *Estimates* | *Conf. Int (95%)* | *P-Value* | *Estimates* | *Conf. Int (95%)* | *P-Value* | *Estimates* | *Conf. Int (95%)* | *P-Value* |
| (Intercept) | -103.73 | -125.92 – -81.54 | **<0.001** | 0.12 | 0.00 – 0.23 | **0.047** | -0.11 | -0.25 – 0.04 | 0.16 | -0.17 | -0.27 – -0.07 | **0.001** | -0.16 | -0.62 – 0.30 | 0.505 |
| Concentration [0.25%] | -6.74 | -42.98 – 29.50 | 0.708 | 0.06 | -0.13 – 0.25 | 0.53 | 0.08 | -0.16 – 0.32 | 0.526 | 0.02 | -0.14 – 0.19 | 0.79 | -0.28 | -1.03 – 0.47 | 0.455 |
| Concentration [0.5%] | 3.52 | -32.85 – 39.89 | 0.845 | 0.1 | -0.09 – 0.29 | 0.276 | 0.14 | -0.10 – 0.39 | 0.23 | 0.07 | -0.09 – 0.24 | 0.382 | 0.02 | -0.73 – 0.78 | 0.949 |
| Concentration [1%] | 29.28 | -6.11 – 64.68 | 0.102 | 0.16 | -0.02 – 0.34 | 0.082 | 0.25 | 0.02 – 0.49 | **0.034** | 0.15 | -0.01 – 0.31 | 0.06 | 0.55 | -0.18 – 1.29 | 0.134 |
| week | 51.38 | 47.44 – 55.31 | **<0.001** | 0.4 | 0.37 – 0.42 | **<0.001** | 0.51 | 0.48 – 0.53 | **<0.001** | 0.33 | 0.31 – 0.34 | **<0.001** | 1.56 | 1.47 – 1.64 | **<0.001** |
| Concentration [0.25%] * week | 3.92 | -2.29 – 10.14 | 0.215 | 0 | -0.03 – 0.04 | 0.834 | 0 | -0.04 – 0.05 | 0.913 | 0.01 | -0.02 – 0.04 | 0.612 | 0.1 | -0.03 – 0.24 | 0.114 |
| Concentration [0.5%] * week | 1.42 | -4.88 – 7.71 | 0.658 | 0 | -0.03 – 0.04 | 0.869 | 0 | -0.05 – 0.04 | 0.864 | 0.01 | -0.02 – 0.04 | 0.7 | 0.04 | -0.09 – 0.17 | 0.537 |
| Concentration [1%] * week | -12.29 | -18.59 – -5.98 | **<0.001** | -0.04 | -0.08 – -0.01 | **0.017** | -0.07 | -0.12 – -0.03 | **0.001** | -0.04 | -0.08 – -0.01 | **0.004** | -0.29 | -0.42 – -0.16 | **<0.001** |
| **Random Effects** | | | | | | | | | | | | | | | |
| $\sigma^2$ | | 2008.71 | | | 0.06 | | | 0.1 | | | 0.05 | | | 0.88 | |
| $\tau_{00}$ | | 303.58 $_{cricket}$ | | | 0.00 $_{cricket}$ | | | 0.01 $_{cricket}$ | | | 0.00 $_{cricket}$ | | | 0.12 $_{cricket}$ | |
| ICC | | | | | 0.05 | | | 0.08 | | | 0.05 | | | 0.12 | |
| N | | 37 $_{cricket}$ | | | 37 $_{cricket}$ | | | 37 $_{cricket}$ | | | 37 $_{cricket}$ | | | 37 $_{cricket}$ | |
| Observations | | 286 | | | 286 | | | 286 | | | 286 | | | 286 | |
| Marginal $R^2$ / Conditional $R^2$ | | 0.871 / NA | | | 0.924 / 0.928 | | | 0.924 / 0.930 | | | 0.916 / 0.920 | | | 0.926 / 0.935 | |

**Table S4:** Results of linear mixed effects model for microfiber concentration vs. growth of the abdomen, thorax, head, and weight of male *G. sigillatus* with cricket identity as random effects and presented with interactions with the cricket age (week). Results considered statistically significant from the control ($p < 0.05$) are presented in bold.

| | Male Microfiber Diet | | | | | | | | | | | | | | |
|---|---|---|---|---|---|---|---|---|---|---|---|---|---|---|---|
| | Weight | | | Head Width | | | Thorax Width | | | Thorax Length | | | Abdomen Length | | |
| *Coeffcient* | *Estimates* | *Conf. Int (95%)* | *P-Value* | *Estimates* | *Conf. Int (95%)* | *P-Value* | *Estimates* | *Conf. Int (95%)* | *P-Value* | *Estimates* | *Conf. Int (95%)* | *P-Value* | *Estimates* | *Conf. Int (95%)* | *P-Value* |
| (Intercept) | -60.62 | -75.70 – -45.53 | **<0.001** | 0.38 | 0.27 – 0.49 | **<0.001** | 0.22 | 0.08 – 0.36 | **0.002** | 0.09 | -0.01 – 0.20 | 0.09 | 0.26 | -0.20 – 0.71 | 0.264 |
| Concentration [0.25%] | -5.18 | -25.29 – 14.94 | 0.606 | -0.11 | -0.26 – 0.04 | 0.138 | -0.13 | -0.31 – 0.06 | 0.17 | -0.09 | -0.23 – 0.05 | 0.18 | -0.2 | -0.81 – 0.40 | 0.506 |
| Concentration [0.5%] | 6.45 | -15.96 – 28.85 | 0.565 | -0.06 | -0.22 – 0.11 | 0.496 | -0.05 | -0.25 – 0.16 | 0.634 | -0.02 | -0.18 – 0.13 | 0.768 | -0.08 | -0.75 – 0.60 | 0.817 |
| Concentration [1%] | 1.59 | -20.95 – 24.13 | 0.888 | -0.05 | -0.22 – 0.11 | 0.531 | -0.03 | -0.24 – 0.17 | 0.749 | -0.05 | -0.20 – 0.11 | 0.545 | 0.01 | -0.67 – 0.68 | 0.988 |
| week | 36.37 | 33.77 – 38.96 | **<0.001** | 0.33 | 0.31 – 0.35 | **<0.001** | 0.43 | 0.40 – 0.45 | **<0.001** | 0.26 | 0.24 – 0.28 | **<0.001** | 1.5 | 1.41 – 1.59 | **<0.001** |
| Concentration [0.25%] * week | 0.77 | -2.61 – 4.14 | 0.655 | 0.01 | -0.02 – 0.04 | 0.427 | 0.02 | -0.02 – 0.05 | 0.362 | 0.02 | -0.01 – 0.04 | 0.252 | 0 | -0.12 – 0.11 | 0.965 |
| Concentration [0.5%] * week | -4.16 | -7.92 – -0.40 | **0.03** | 0 | -0.03 – 0.03 | 0.944 | -0.01 | -0.05 – 0.03 | 0.666 | -0.01 | -0.04 – 0.02 | 0.633 | -0.07 | -0.20 – 0.05 | 0.259 |
| Concentration [1%] * week | -2.37 | -6.22 – 1.48 | 0.227 | 0.01 | -0.02 – 0.04 | 0.545 | 0 | -0.04 – 0.04 | 0.999 | 0.01 | -0.02 – 0.04 | 0.562 | -0.07 | -0.20 – 0.06 | 0.299 |
| **Random Effects** | | | | | | | | | | | | | | | |
| $\sigma^2$ | | 803.27 | | | 0.06 | | | 0.08 | | | 0.05 | | | 0.91 | |
| $\tau_{00}$ | | 158.88 $_{cricket}$ | | | 0.00 $_{cricket}$ | | | 0.00 $_{cricket}$ | | | 0.00 $_{cricket}$ | | | 0.03 $_{cricket}$ | |
| ICC | | | | | 0.01 | | | 0.04 | | | 0 | | | 0.03 | |
| N | | 47 $_{cricket}$ | | | 47 $_{cricket}$ | | | 47 $_{cricket}$ | | | 47 $_{cricket}$ | | | 47 $_{cricket}$ | |
| Observations | | 372 | | | 372 | | | 372 | | | 372 | | | 372 | |
| Marginal $R^2$ / Conditional $R^2$ | | 0.891 / NA | | | 0.915 / 0.916 | | | 0.918 / 0.922 | | | 0.875 / 0.875 | | | 0.923 / 0.926 | |
